# Physicochemical and microbiological profile of the potable water of Eastern Himalayan State Sikkim: An Indication of Severe Fecal Contamination and Immediate Health Risk

**DOI:** 10.1101/508283

**Authors:** Ashish Kumar Singh, Saurav Das, Samer Singh, Nilu Pradhan, Santosh Kumar, Varsha Rani Gajamer, Yangchen D. Lepcha, H. K. Tiwari

**Author notes:** Corresponding Author: Dr. Hare Krishna Tiwari, Department of Microbiology, School of Life Sciences, Sikkim University, Gangtok, Sikkim. Email ID. Designed the study as well as supervised the study (H. K. Tiwari). Performed the experiment (Ashish Kumar Singh). Analyzed the data and prepared the manuscript (Saurav Das, Samer Singh). Helped in the sample collection, survey and lab experiments (Nilu Pradhan, Varsha Rani Gajamer). Helped in manuscript preparation (Santosh Kumar). Helped in the selection of study area and official documentation regarding the study (Yangchen D. Lepcha).

## Abstract

Increasing population, rapid urbanization and climate change have immensely affected the freshwater sources around the world. The continuous decline in the number of natural potable water sources raises serious concern about the overall health of the human population. Developing countries are the most affected in this regard due to lack of proper hygienic maintenance protocols. Sikkim an Eastern Himalayan state with mountains as predominant topological features, harbors several natural spring water (SW). These spring waters are the primary source of potable water for the population in the four districts of the state viz. East (E), West (W), South (S) and North (N). Several incidences of water-borne diseases and the relative lack of scientific evaluation reports on the water quality of the area have educed this study. Lack of any standard filtration and purification practice among the population is one of the prime factors for the outspread of different waterborne pathogen in the state. The people of the state mostly use boiling as a dominant method of water purification, while only a small percentage of people in the West district were found to use modern standard purification system (W = 30%). The rainy season was found to be the major contributor of different diseases (E = 86%; W=100%; S=100%; N=80%) and statistical analysis of the fecal coliforms of the different season also indicated a significant difference at p < 0.05. There was no statistical significance among the physicochemical parameter of the SW but surprisingly the water from the four districts was recorded with traces of highly toxic heavy metals like mercury/WHO limit (0.001-0.007mg/l/0.001) as well as lead/WHO limit (0.001-0.007mg/l/0.05) and selenium/WHO limit (0.526-0.644 mg/l/0.01) which was above the WHO permissible limit. Piper analysis showed that water was dominated by cation sodium ion (Na^+^) and bicarbonate (HCO_3_^-^) anion and the water can be categorized as Mg-HCO_3_^-^ type. Pairwise Pearson correlation showed a significant correlation between Electrical conductivity and TDS (r = 0.998/1.00) as well as alkalinity and turbidity (0.993/1.00). The microbial confirmatory test showed severity in fecal contamination with high counts of Total Coliform (TC), *Escherichia coli* (EC) and *Enterococcus* (EN). Highest TC was recorded from W (37.26/ml) and lowest in N (22.13/ ml) in spring water. Highest contamination of *Escherichia coli* and *Enterococcus* was found in E (EC = 8.7/ml; EN = 2.08/ml) followed by S (EC = 8.4/ml and EN = 2.05/ml). It was found that community reservoir (CR) tank was more contaminated than SP, which indicates the negligence in maintenance and fecal contamination during transportation to the reservoir. Though household water was least contaminated compared to CR and SP, but it fails in WHO standard criteria for drinking water. These results indicate an immediate health risk of the resident of the state and which needs to be taken care of sooner as possible by protecting the important potable sources with required policies and regulations.

## 1. INTRODUCTION

Urbanization and increasing anthropogenic activity have wedged the earth’s natural environment and negatively impacted human health (Fronczyk et al., 2016). Contamination of water sources has become a concern for the scientific community as it is directly related to the health index of people. Around the world, water contamination has led to several widespread illnesses like diarrhea, dysentery, cholera, typhoid and death on a regular basis. Developing countries are on the alarming list due to insufficient availability of pure potable water and lack of good quality of health care (World Health Organisation, 2018). Bacterial diseases are the most common when the population comes to contact with contaminated water and it is also becoming tougher to treat due to the increasing antibiotic resistance among the pathogens. These diseases can be fatal and life-threatening depending on the host susceptibility. Thus, access to safe drinking water is critical to human health and development (Pruss et al., 2002). Rural remote areas of Sikkim, India still depend on using natural water as a potable source without proper filtration and treatments. Three main drinking water chains are known to play a significant role in the safety of drinking water: the quality of raw water at the source, the purification process of water and the distribution system of water (Ikonen et al., 2017). These all three chain system affect the physicochemical characteristics as well as microbial composition (Geldreich, 1989). Several microbial communities survive in the water system and in a favorable environment, they start multiplying in the water and depreciate the water quality (Ikonen et al., 2017). Diarrhea has become the fourth leading cause of death worldwide and in 2015, it caused an estimated 1.3 million death of children under the age of 5 years (Troeger et al., 2017).

The present global population is around 7.6 billion and growing at the rate of 83 million per year, globally (United Nations New York, 2017). In fact, of the projected population growth to 8.6 billion by 2030 and some 8.9 billion by 2050, almost all will be in low-income countries (World Health Organisation, 2018). The increase in population leads in rising competition for freshwater between agriculture, urban and industrial uses that impact pressure on rural and urban areas (UN-Water, 2006). Surface water and groundwater act as the major drinking water source to the entire human community and other organisms (Sharma et al., 2017). Groundwater is considered a reliable source of fresh water which is easy to access for various purposes such as domestic, industrial, irrigation etc. (Kumar et al., 2017). Worldwide, approximately 1.5 billion peoples are directly or indirectly dependent on the groundwater for their domestic and agricultural needs (Mukherjee and Singh, 2012). The geochemical process like dissolution, hydrolysis, precipitation, adsorption and ion exchange, as well as oxidation-reduction and biochemical reaction, are major controlling factors for the chemistry of groundwater. It is quite interesting, about 80% rural and 50 % urban people of India directly depends on the groundwater whereas 56 % rural Indian access potable water from tube wells, 14% from open wells and 25% by supplied water system (Kumar et al., 2017). One of the reports mentioned that household which is getting drinking water from ‘improved’ sources also showed that 42% in urban and 60% in rural households were, in reality, receiving contaminated water (WaterAid, 2017). In India, most of the sewage finds its way into the city drain and finally into the rivers contributing to the poor quality of water sourced from the river. Due to unavailability of proper fecal sludge treatment, most of the sewage and fecal sludge find own way to the ground and that cause increased the frequency of groundwater pollution of bacteriological contamination (WaterAid, 2017). According to the survey report of lancet 2014, about 24% of children in urban areas and 55% of children in the rural area are falling ill due to contaminated water (Johri et al., 2014). They also reported that about 11% of urban households and 23% of rural households have suffered infants death (Johri et al., 2014).

Sikkim (270 05’ to 280 07’ N latitudes and 870 59’ to 880 56’E longitudes), an Eastern Himalayan state of India, lies between Nepal and Bhutan (Tambe et al., 2012b). The state is categorized by a tremendous scarcity of potable and treated drinking water for the rural as well as urban population. The crisis is much more severe in the urban and city areas relatively, because of deficiency of sources of fresh water and various types of contamination of water in and around these areas. Though the Himalayan range is a source of countless perennial rivers, and the mountain people depend largely on spring water for their sustenance. The mountain springs, locally known as *“Dharas”*, are the natural discharges of groundwater from various aquifers. Some of the springs are considered sacred, are revered as Devithans and are protected from biotic interferences (Tambe et al., 2012b). In rural areas of Sikkim, a major source of water for drinking and domestic purpose is supplied by spring water. About 80% of the total rural community is solely dependent on spring water for their domestic and drinking purpose (Tambe et al., 2012a). The use of untreated surface water sources for drinking and domestic purpose remains a major threat to public health, as these could serve as a reservoir for the transfer of waterborne pathogens (Tambe et al., 2012b). The dependence of the people of the state on spring water solely defines the importance of this study. This study reports the water quality of springs water, community reservoir and household water of four districts *viz*. East (E), West (W), South (S) and West (W), in terms of both physicochemical and microbiological qualitative and quantitative index. Lack of such previous reports makes it an important foundation to structure and device the future water treatment protocols and to formulate government policies to tackle the immediate health risks.

## 2. Material and Methods

### 2.1 Description of Study Site

The study was conducted in different villages of Sikkim, which is a NorthEastern state of India. The State is located off the slopes of Eastern Himalayas between the latitude of 27° 05” and 28° 07” North and the longitudes of 88° 28” and 88° 55” East, covering approximately 115 km from North to South and 65 km from East to West. The landscape of this area varies with an altitude between 300 to 8583 meter above sea level that comprises lower, middle and higher hills, alpine zones and snowbound land. Sikkim is a multi-ethnic state in which population is divided into tribal and non-tribal groups (Tambe et al., 2012a). The Himalayan range is one of the important sources of numerous permanent rivers, though the Himalayan range is a source of countless perennial rivers and springs. The mountain peoples are mostly dependent on the spring water for their drinking and household purpose. The springs are the natural release of groundwater from different aquifers which are locally known as ‘Dharas’. Approximately, 80% of the rural households in Sikkim depend on spring water for their daily water needs. Among the springs, some springs are considered as blessed, are known as ‘Devithans’ and confined from any biotic interferences (Tambe et al., 2012a). There are several household levels, community, and village levels storage tank were prepared by Rural Management and Development Department (RMDD), Government of Sikkim in which spring water is collected to be utilized by the community for household purpose and up to some extent for irrigation of kitchen gardens as well as greenhouse crops (Arrawatia and Tambe, 2012).

**Fig. 1:**
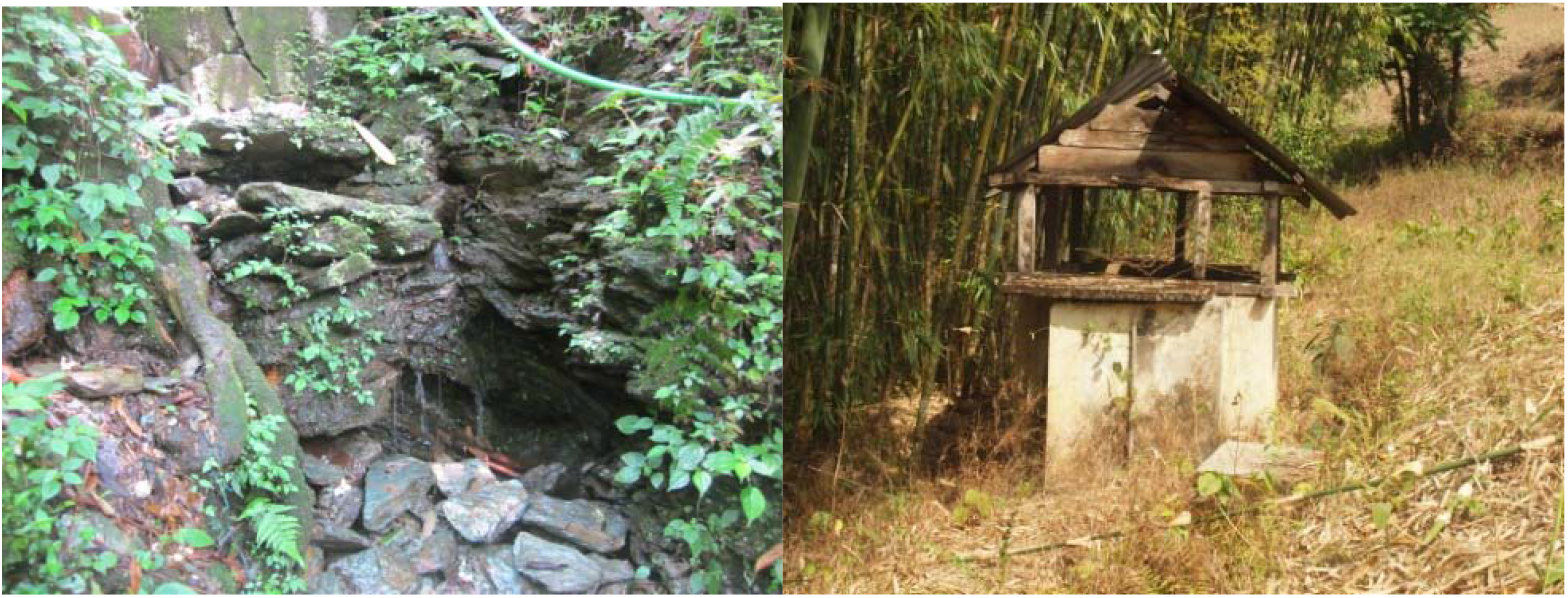
Photograph of the study areas (a) Spring water (b) A typical community reservoir – where water from the springs was stored before being distributed to the households (c) A Sikkimese family standing in front of a newly constructed community reservoir.

**Fig. 2:**
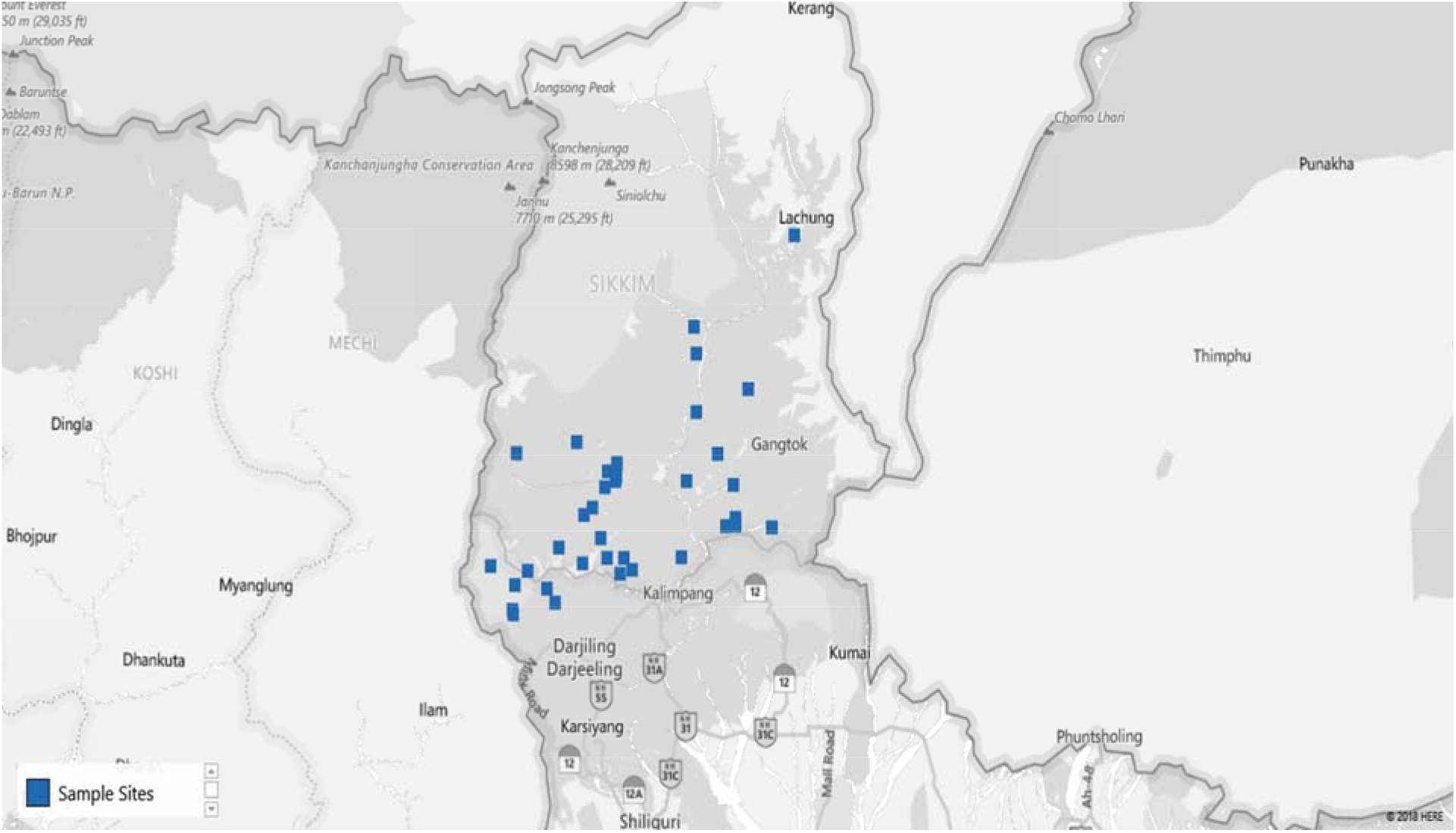
Map of the study site (blue dots specify the villages from where samples were collected). The map was collected from the: (Spring Detail: http://www.Sikkimsprings.org/dv/view.php) [Map chart was prepared in Microsoft Excel (office 365)] (Supplementary File: Table S1).

### 2.2 Study Design

For current study, 40 springs that served as the potable water source and 40 community reservoirs were selected (10 from each district: East, West, South and North district). For household water samples, 10 villages were selected from each district with a total of 40 villages and water samples from 5 households of each village were collected (10 x 4 x 5 = 200 samples). Springs were selected based on the latitude, the number of households depended, and perennial nature (**Supplementary Table. S1**). Different Gram Panchayat Unit (GPU) or village level blocks were selected on the basis recent reports of water-borne diseases incidence in villages of that GPU. At each household, respondent was requested to provide water which they are using for drinking. A structured questionnaire was used for the survey (**Supplementary file: S2**)

**Fig. 3:**
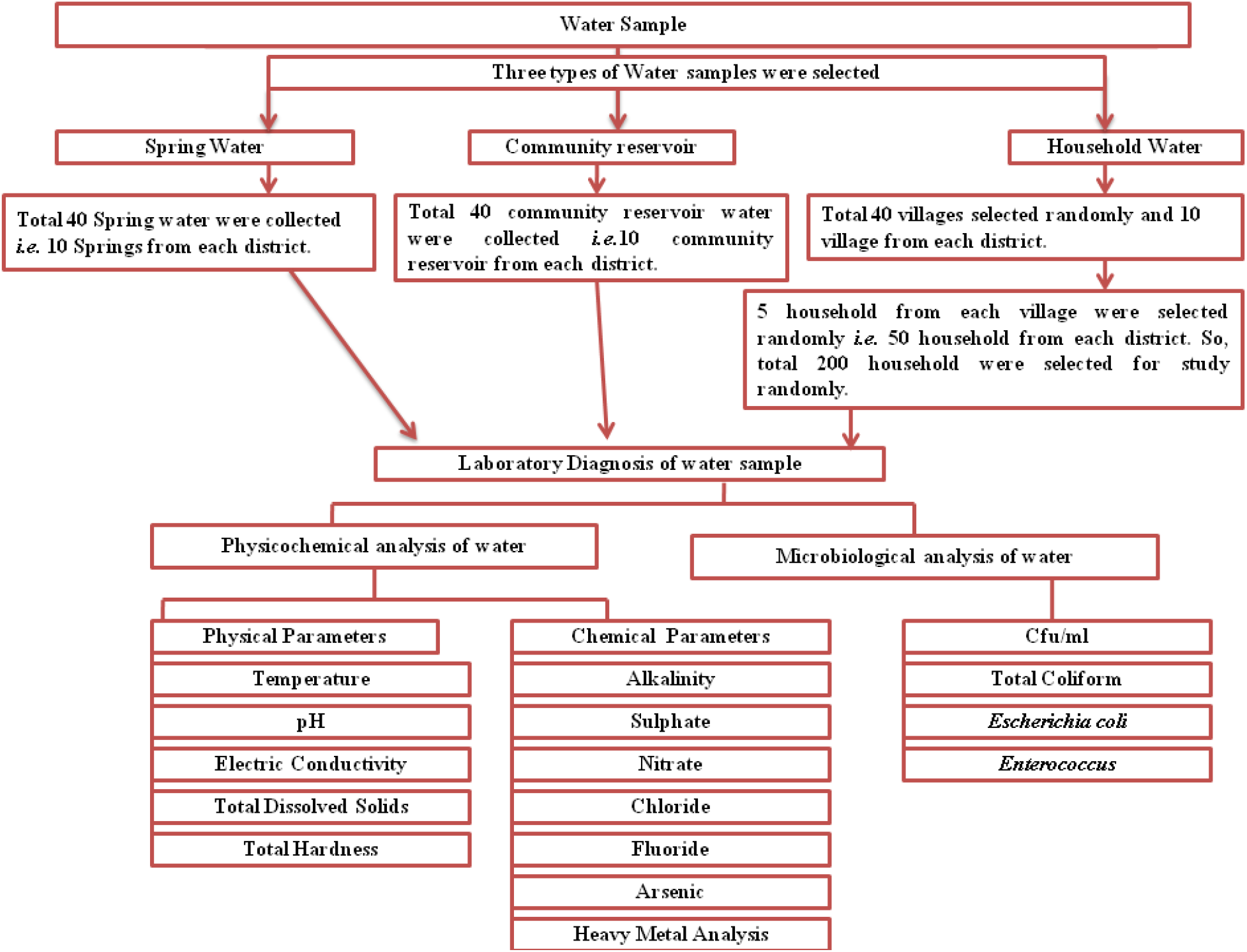
Flowchart of the study design. Three different types of water samples were collected: (i) Spring water; (ii) Community reservoir water and (iii) household water and evaluated for the microbial and physicochemical analysis as indicated.

### 2.3 Sampling procedure

Water samples were collected in 1L sterile, wide mouth, plastic bottles (Nalgene, USA). Before collection, containers were sterilized by autoclave and washed with aqueous sodium thiosulfate solution [100 g/l (w/v)]. While collecting the sample, the bottles were completely submerged into the water and it was opened inside the water source to avoid the air contamination. The container was filled by holding it diagonally; lower part submerged up to 30 cm with the mouth facing slightly upwards. A gap of 2 cm was left between the cap and water to provide sufficient airspace for mixing of water before analysis (Gillet et al., 2009).

### 2.4 Physicochemical analysis

Physicochemical parameters of spring water like Temperature (°C), pH, Electrical Conductivity (μS/cm), Total Dissolved Solids (TDS) (mg/l) was measured by multi-probe potable meter (Hi-media, Mumbai, India). Alkalinity (ppm), Total Hardness (TH) (mg/l), Sulphate (mg/l), Nitrate (mg/l), Chloride (mg/l), Fluoride (mg/l) were measured by Aqua check water testing kit provided by Hi-media, Mumbai, India. The elemental detection was performed by IC-PMS (Inductive Coupled-Plasma Mass Spectroscopy) (Perkin-Elmer NexIon 300X, USA). The detected elements were, In (Indium), Ba (Barium), Pb (Lead), Ag (Silver), Al (Aluminum), As (Arsenic), Ba^-1^ (Barium), Be (Beryllium), Bi (Bismuth), Ca (Calcium), Cd (Cadmium), Co (Cobalt), Cr (Chromium), Cs (Cesium), Cu (Copper), Fe (Iron), Ga (Gallium), S (Sulphur), K (Potassium), Li (Lithium), Mg (Magnesium), Mn (Manganese), Na (Sodium), Ni (Nickel), Mo (Molybdenum), R (Rubidium), Se (Selenium), Sr (Strontium), Ti (Titanium), U (Uranium), B (Boron), V (Vanadium), Zn (Zinc), Hg (Mercury), Si (Silicon), P (Phosphorus), N (Nitrogen), Cl (Chlorine), Zr (Zirconium), Xe (Xenon), Sn (Tin), I (Iodine), Ce (Cerium). All the water samples were collected and analyzed directly without any pretreatment except for Arsenic and Nitrate. Water samples for analysis of arsenic and nitrate were mixed with the acid (1M HCl in the 5ml for 500 water sample) (Fisher et al., 2015). The physicochemical parameter of community water and household water were not measured as they are just replicated source from the springs (spring water is collected to community reservoirs tank which was supplied to the household). The initial physicochemical test of community reservoir and household water showed a similar chemical profile.

### 2.5 Microbiological Analysis

For Microbiological analysis water samples were collected separately in 1-liter sterile containers. Samples for microbiological analysis were transported in ice cooled condition. Water samples were collected in triplicate. Membrane Filtration technique was used for the detection of *E. coli* and Total Coliform counts (Collee, J. G., Miles, R. S. and Wan, 2014). For detection of the total coliform, membrane filter was placed on LES-Endo agar and the bacterial colony was counted by Colony Counter (Cole-Parmer, India). For detection of *E. coli*, membrane filters were cultured on Rapid Hi-chrome Agar (Hi-media, M1465). Colonies with blue-green color with fluorescence under UV exposure and indole positive tests were identified as *E. coli*. All plates were incubated at 35 ± 2°C for 24 hours (h). For detection of *Enterococcus* group, the filter was incubated on m-*Enterococcus* agar at 41°C for 48 h (American Public Health Association, 1999). Coliform and fecal streptococci density were calculated using the following equation as described by American Public Health Association, 1999 (American Public Health Association, 1999).

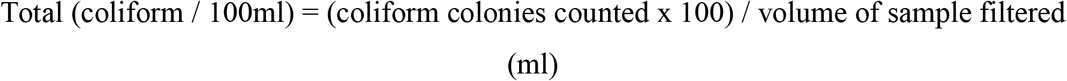

### 2.6 Statistical Analysis

Results were analyzed using Microsoft Excel (Office 365). Wilcoxon signed rank test was performed to compare the log concentration of *E. coli, Total coliform*, and *Enterococci* between Springwater, Community reservoir, and Household water samples. One Way ANOVA was used to compare colony forming unit in different water sources and significance of the difference in the physicochemical parameters. The piper plot was used to categorize the water based on total hardness. The piper plot was constructed using AquaChem v 5.0. Pearson correlation was used to determine the correlation between the different physicochemical parameters. The correlation coefficient (r) was calculated using Microsoft Excel (office 365) and correlation plots were prepared using R studio (package: corrplot and PerformanceAnalytics).

## 3.0 Results

### 3.1 Microbiological Water quality

#### 3.1.1 Spring Water

All the spring water (100%,) in the month of July-August (rainy season) comes under the category of intermediate risk level. Whereas, in the month of November-December, East Sikkim springs showed 80% intermediate level and 20% low-risk level for the total coliform count. In the same months November-December, the South Sikkim water samples showed 70% intermediate and 30% at the low-risk level of contamination. Whereas spring waters from the North Sikkim showed 50% intermediate level and 50% low-risk level for total coliform concentration (**Fig.4**). Water samples from West Sikkim had come under intermediate risk level in the all the three seasons. It was also found that about highest number (90%,) of spring water of East Sikkim was contaminated *E. coli* followed by South Sikkim (80%,) and West Sikkim (60%,) and North showed least contamination (50%,) (**Fig.5**). The results showed that all the spring water samples were highly contaminated in the month of July-August in all the four districts and least contaminated in the month of March-April.

**Fig. 4:**
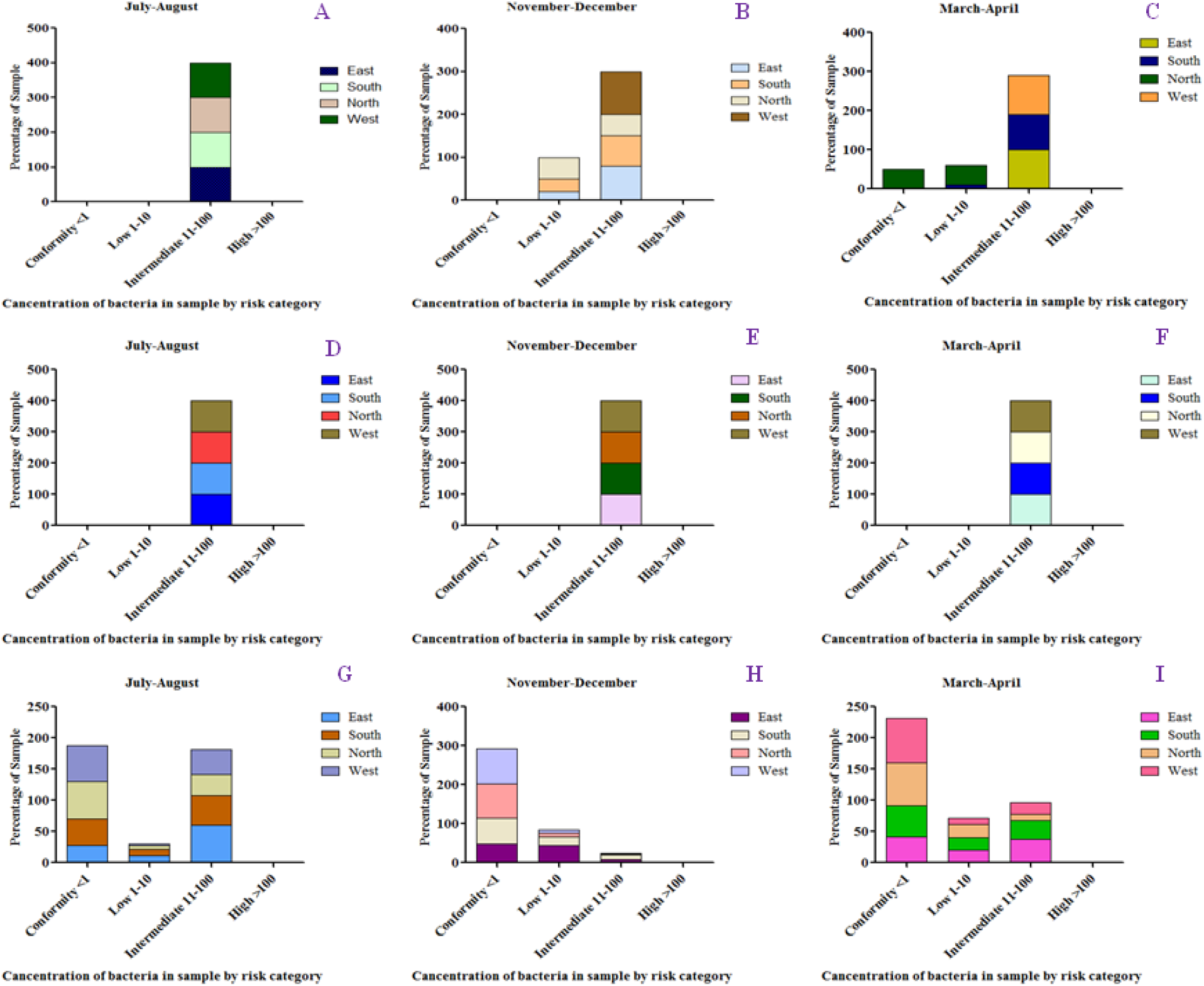
a Characterization of water samples by risk category on the basis of bacterial concentration. A-C: Total coliform concentration Spring Water, D-F: Total coliform concentration in community reservoir, G-H: Total coliform concentration in the community household water in the different season.

**Fig. 5:**
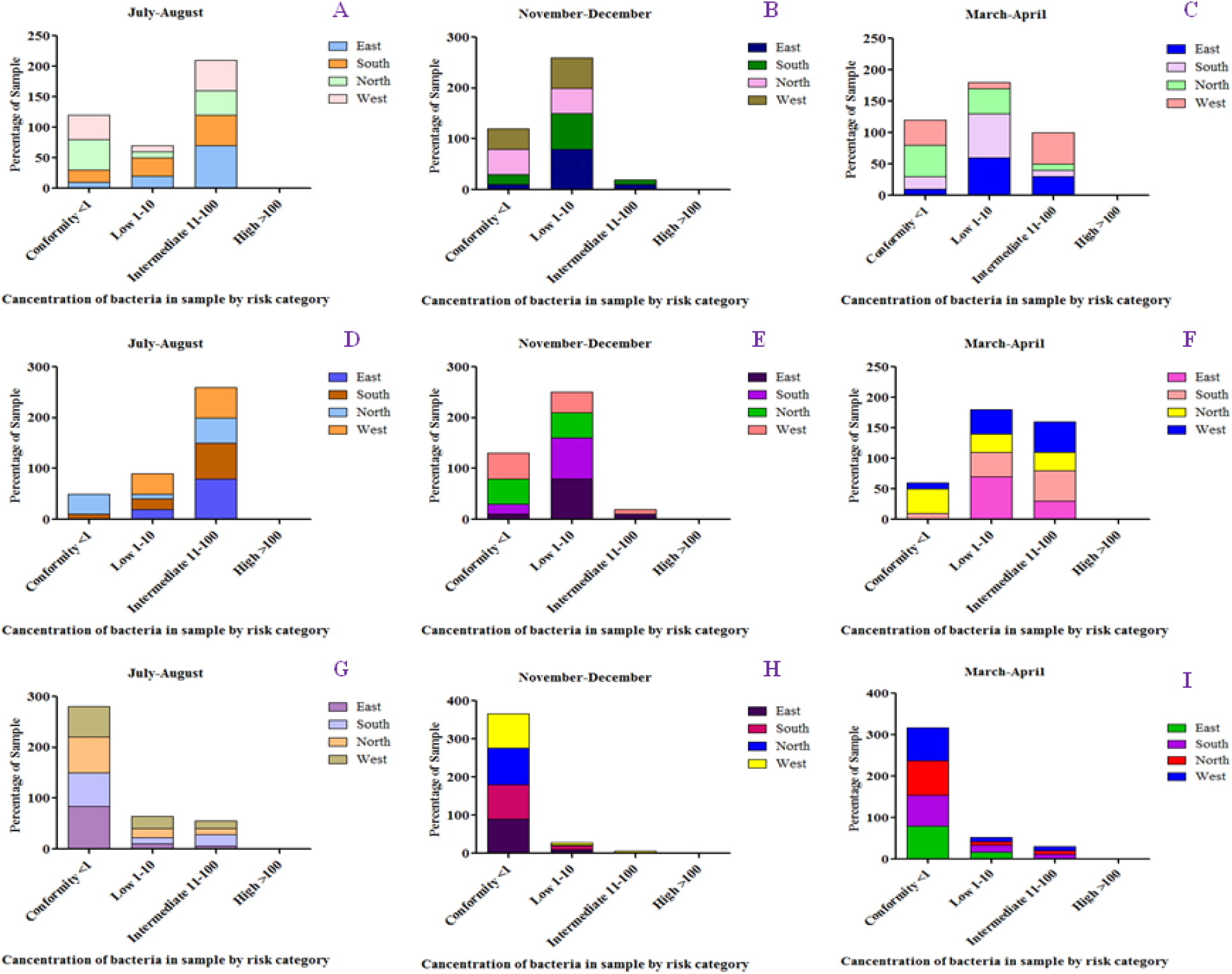
Characterization of water samples by risk category on the basis of bacterial concentration. A-C: *E. coli* concentration Spring Water, D-F: *E. coli* concentration in community reservoir, G-I: *E. coli* concentration in the community household water in the different season.

#### 3.1.2 Community reservoir Tank

It was found that all the community reservoirs contained more than > 10 <100 total coliform count and comes under the category of intermediate risk level (**Fig.4**). Based on *E. coli* detection, about 79% of community reservoir was found to be contaminated with *E. coli*. The highly (90%) *E. coli* contamination was reported in East Sikkim whereas least contamination (56%) was present in North Sikkim (**Fig.5**).

#### 3.1.3 Household Water

It was surprising to observe that besides the SW and CR, HW samples were also contaminated with total coliform above the permissible value, although few of the households used boiling before consumption. High concentration of total coliform was observed in the East Sikkim household whereas least in the West Sikkim (26%) (**Fig.4**). But in the case of *E. coli* contamination, it was observed that West and South Sikkim household water samples were highly (23%) contaminated whereas East showed least (15.33%) contamination (**Fig.5**). As compared to SW and CR, most of the household water comes under the conformity to low risk level and very less household water samples had come under the intermediate level which was mostly in the month of July-August (rainy season).

### 3.2 Household water treatment profile

During the study, we collected 200 household water samples (10 villages x 4 district x 5 households from each district = 200 water samples) and discovered most of the households never used any kind of standardized treatment or filtration process prior to taking the water as a potable source. Boiling is the only or major source of decontamination used by the household in four districts of the state (E: 66%; W: 60%; N: 80%; S = 56%), only a few percentages of the people used filtration prior to consumption or using it for other purposes (E: 14%; W:30%; S:14% and N: 4%). While, still a major portion of the population consumes the raw water directly from the source (spring water) (E: 20%; W: 10%, S: 30% and N: 16%) (**Fig. 6**).

**Fig. 6:**
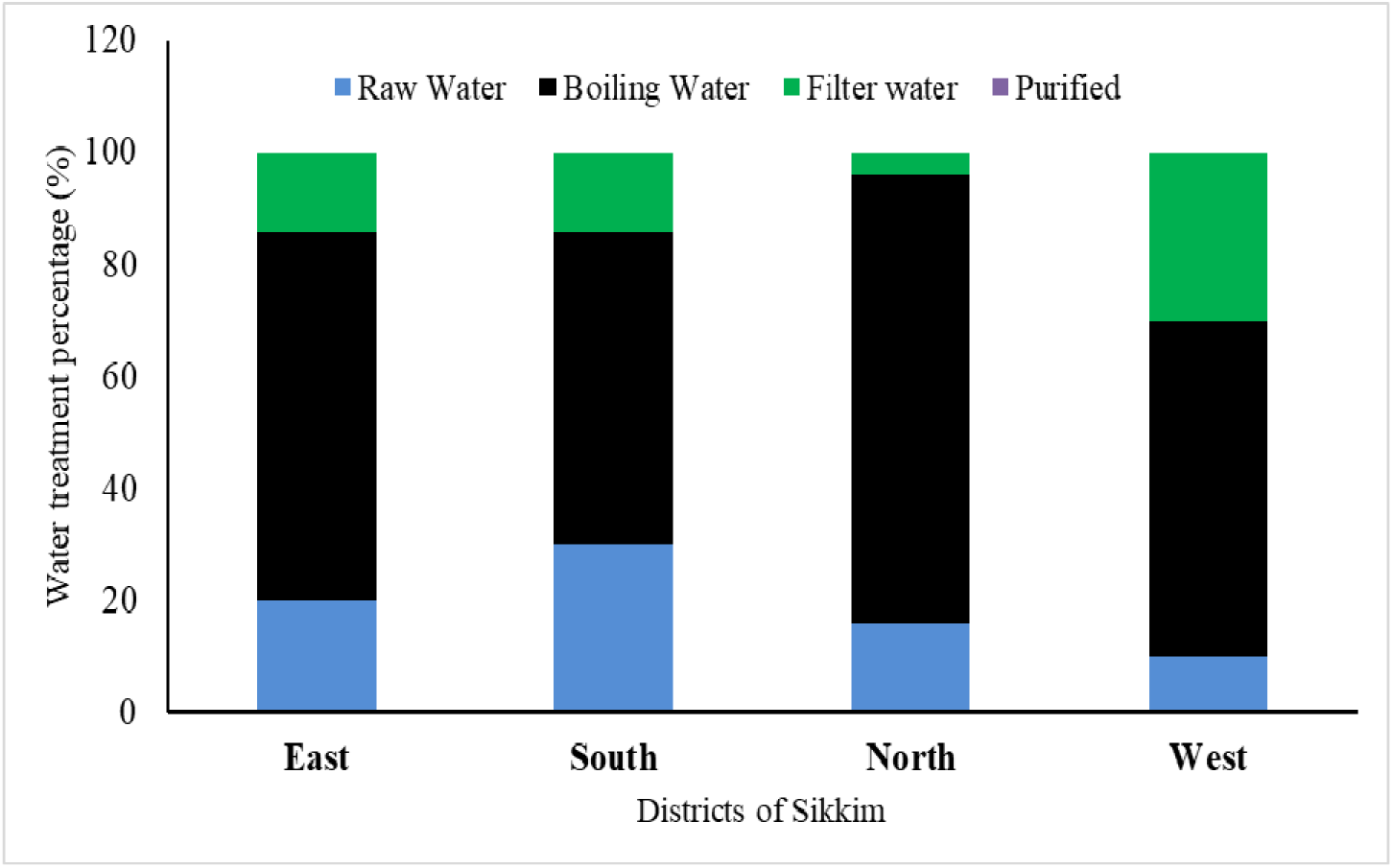
Different water treatment profile among the households of different districts of Sikkim. Boiling is the most common method used by residents prior to consumption.

However, the household water with prior boiling was also recorded with TC and E. coli (EC) count above the permissible guideline. Both, in the case of TC and EC, all the household samples which were boiled prior consumption was found to be in the range of intermediate to low risk category. Rainy season or July – August (JA), month showed the highest contamination. East and South district showed the highest contamination compared to the West and North district (**Fig. 7**).

**Fig. 7:**
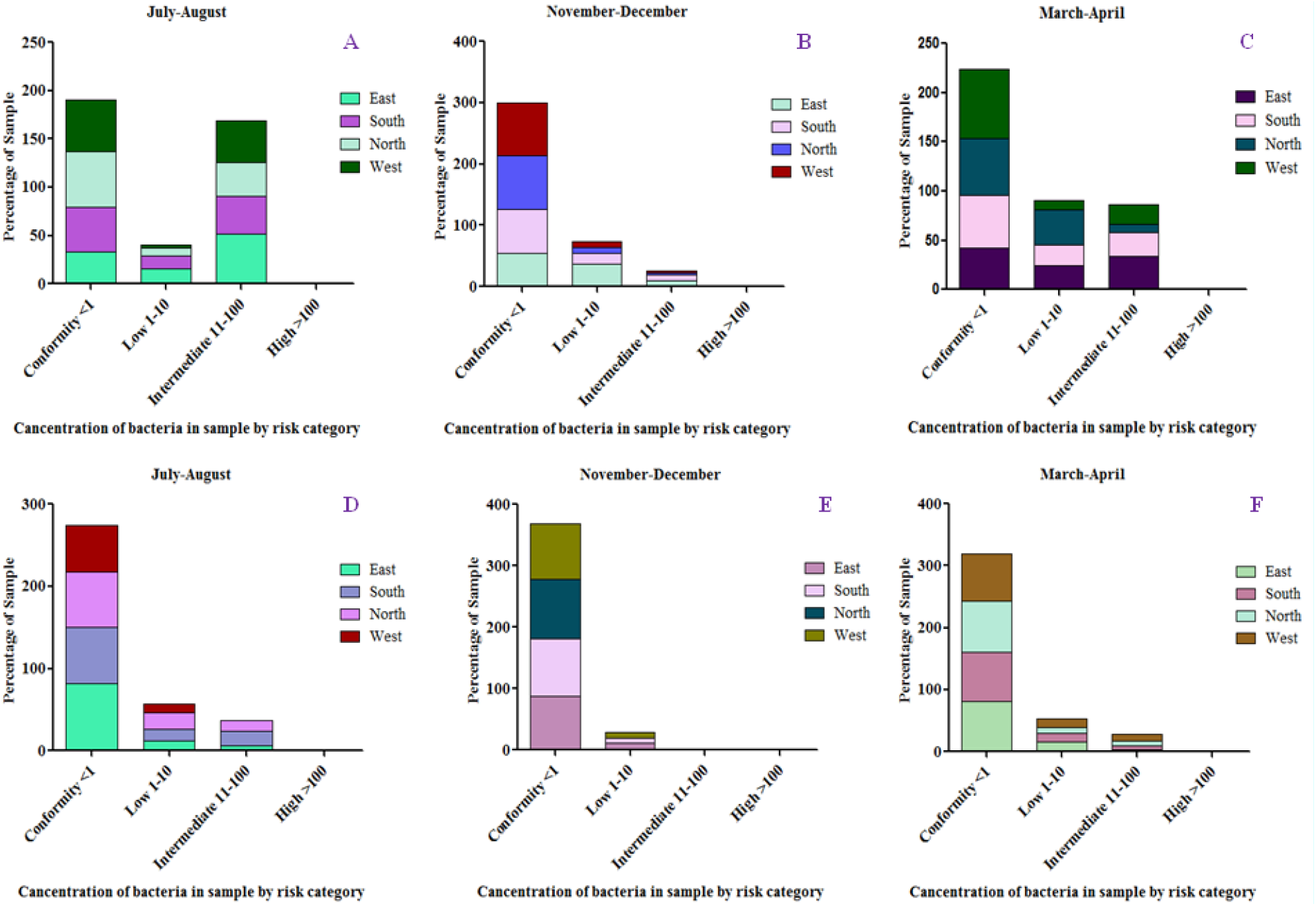
Characterization of water samples by risk category based on bacterial concentration. A-C: Total coliform concentration of household water, D-F: *E. coli* concentration in Household water of that water samples which used boiling methods for treatment.

In the survey, respondent declares that the rainy season was responsible for most of the water-borne diseases. Calculating the percentage of disease incidence, based on the response, it was found that 86% of the disease recorded from the East district had occurred in the rainy season in comparison to the 14% recorded disease in summer. The rainy season was 100% responsible for diseases from South and West district. While 80% of the disease recorded from North was in the rainy season and 3% in the spring season (**Fig. 8**).

**Fig. 8:**
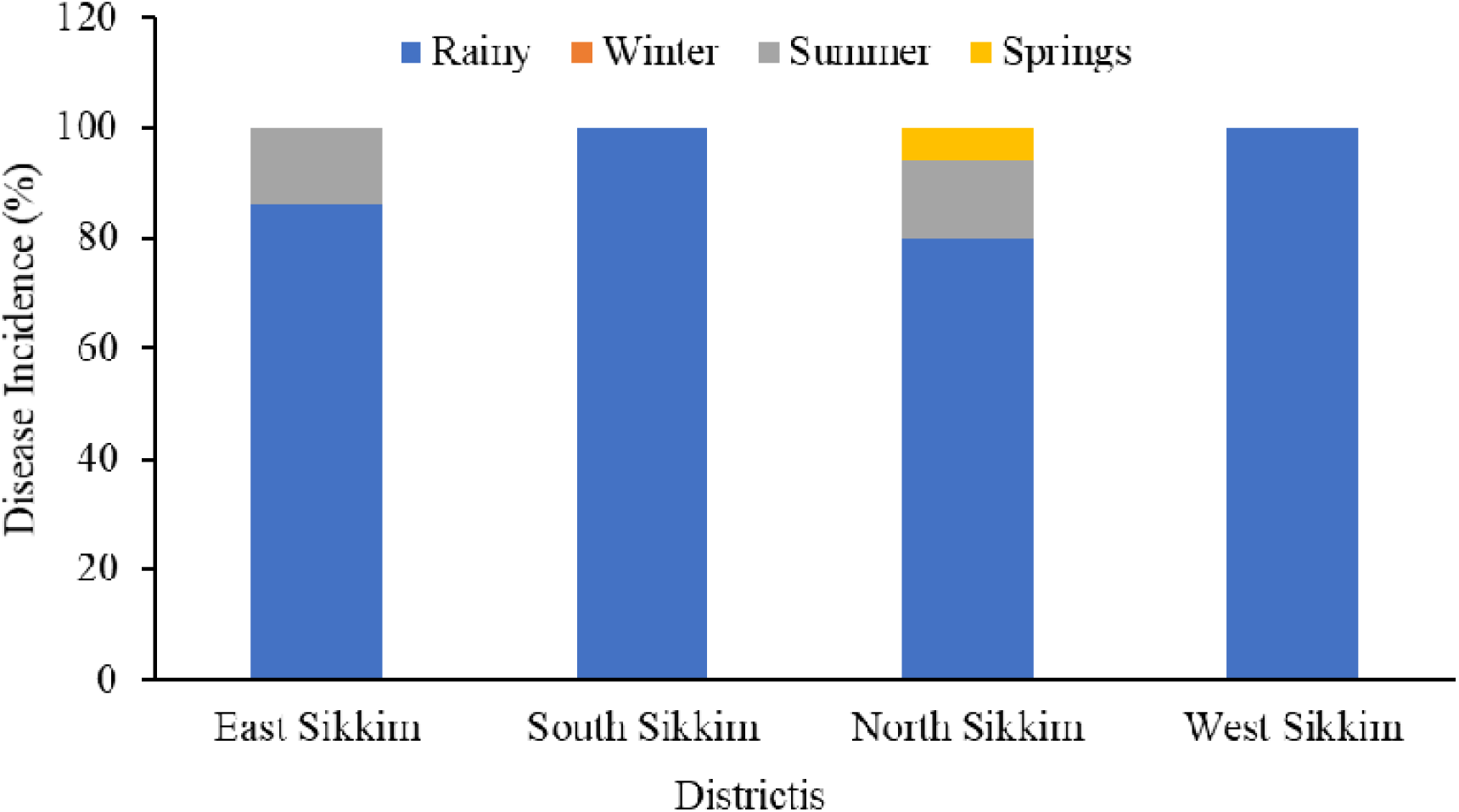
The occurrence (percentage) of disease based on community response. The most disease prone time in Sikkim was rainy season3.3 Physicochemical Parameter

The concentration of total dissolved solids (TDS) in the four districts varied widely from 13.5 to 41.5 mg/l, with East district having the highest TDS of 41.5 mg/l while the having lowest in North (13.5). The total hardness ranged from 21-37 mg/l, with West district having the highest concentration of 37 mg/l whiles the lowest in North (21). All the waters were near neutral in nature within pH range of 6.52 - 7.18, which was higher than standard guidelines like WHO. The temperature of the water ranged between 18.6 – 20.8°C. Turbidity recorded was in the range of 1.5 −2.6 NTU. There was no significant difference between the physical parameters at p <0.05 except the Electrical conductivity (EC) that ranged between 96 – 188.1 μS/cm, the highest value of 188.1 μS/cm was recorded from the South district and the lowest value of lowest 96 μS/cm was recorded from the North district (**Table. 1**).

**Table. 1:**
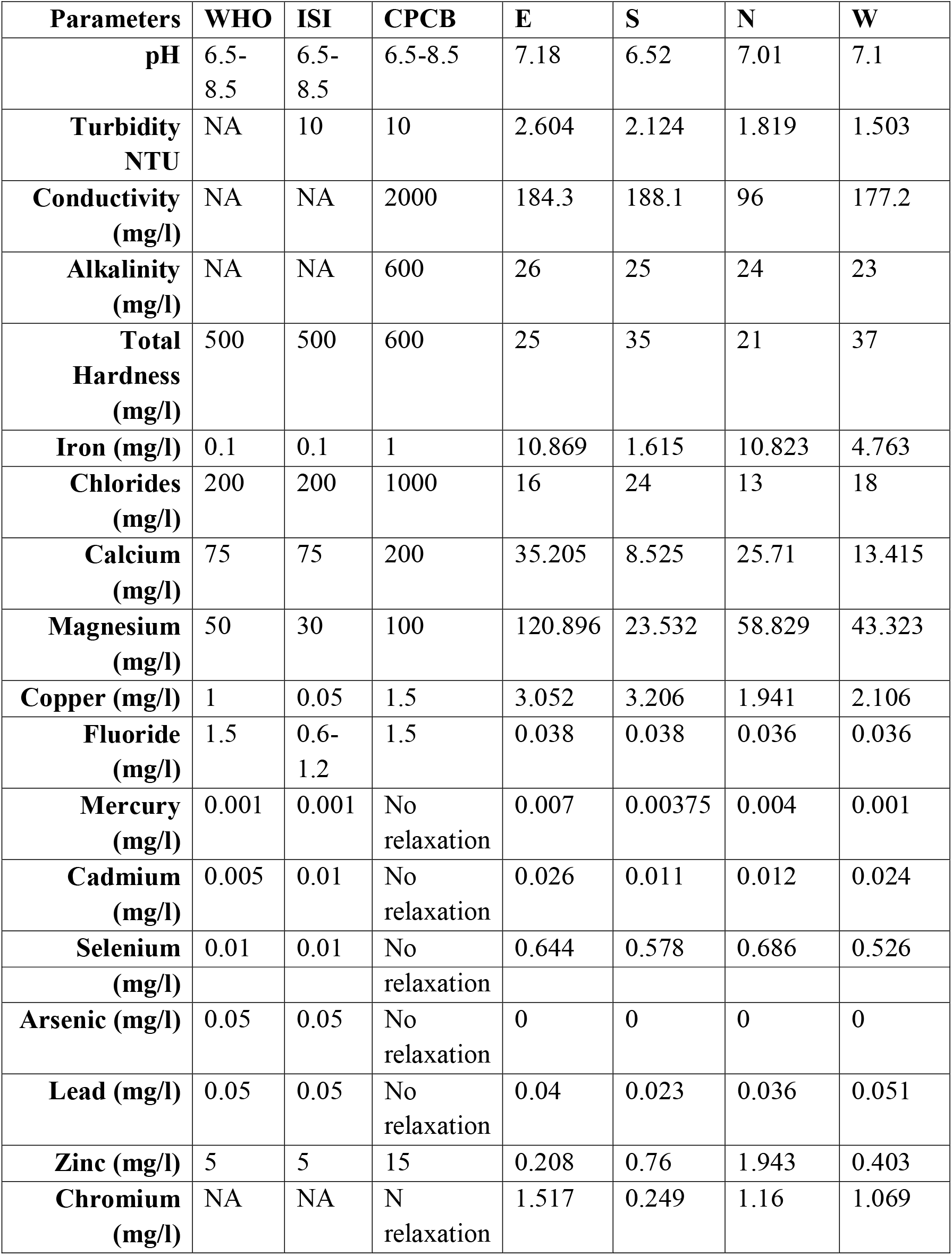
Comparative physicochemical parameter with the standard guidelines of WHO (2017, ISI (2012), and CPCB, (2017).

| **Parameters** | **WHO** | **ISI** | **CPCB** | **E** | **S** | **N** | **W** |
| --- | --- | --- | --- | --- | --- | --- | --- |
| **pH** | 6.5-8.5 | 6.5-8.5 | 6.5-8.5 | 7.18 | 6.52 | 7.01 | 7.1 |
| **Turbidity NTU** | NA | 10 | 10 | 2.604 | 2.124 | 1.819 | 1.503 |
| **Conductivity (mg/l)** | NA | NA | 2000 | 184.3 | 188.1 | 96 | 177.2 |
| **Alkalinity (mg/l)** | NA | NA | 600 | 26 | 25 | 24 | 23 |
| **Total Hardness (mg/l)** | 500 | 500 | 600 | 25 | 35 | 21 | 37 |
| **Iron (mg/l)** | 0.1 | 0.1 | 1 | 10.869 | 1.615 | 10.823 | 4.763 |
| **Chlorides (mg/l)** | 200 | 200 | 1000 | 16 | 24 | 13 | 18 |
| **Calcium (mg/l)** | 75 | 75 | 200 | 35.205 | 8.525 | 25.71 | 13.415 |
| **Magnesium (mg/l)** | 50 | 30 | 100 | 120.896 | 23.532 | 58.829 | 43.323 |
| **Copper (mg/l)** | 1 | 0.05 | 1.5 | 3.052 | 3.206 | 1.941 | 2.106 |
| **Fluoride (mg/l)** | 1.5 | 0.6-1.2 | 1.5 | 0.038 | 0.038 | 0.036 | 0.036 |
| **Mercury (mg/l)** | 0.001 | 0.001 | No relaxation | 0.007 | 0.00375 | 0.004 | 0.001 |
| **Cadmium (mg/l)** | 0.005 | 0.01 | No relaxation | 0.026 | 0.011 | 0.012 | 0.024 |
| **Selenium** | 0.01 | 0.01 | No | 0.644 | 0.578 | 0.686 | 0.526 |
| (mg/l) |  |  | relaxation |  |  |  |  |
| **Arsenic (mg/l)** | 0.05 | 0.05 | No relaxation | 0 | 0 | 0 | 0 |
| **Lead (mg/l)** | 0.05 | 0.05 | No relaxation | 0.04 | 0.023 | 0.036 | 0.051 |
| **Zinc (mg/l)** | 5 | 5 | 15 | 0.208 | 0.76 | 1.943 | 0.403 |
| **Chromium (mg/l)** | NA | NA | N relaxation | 1.517 | 0.249 | 1.16 | 1.069 |

On the piper diagram (Piper, 1944), compositional distribution of chemical showed that the water from the four districts was dominated with cation Na^+^ and which did not significantly varied as a function of sample location, whereas anion composition was overall dominated by bicarbonate (HCO_3_^-^) ions. Based on the piper analysis water can be classified as Mg-HCO_3_^-^ type and categorized as shallow fresh groundwater (**Fig. 9**). The water chemistry analysis showed that the water samples were rich in magnesium and iron with a range of 43.32 – 120.896 mg/l and 1.615-10.869 mg/l respectively. Copper was also detected in significant concentration from 1.94 – 3.05 mg/l. The water samples were also recorded with traces of mercury, selenium, and chromium in the range of 0.001 – 0.007 mg/l, 0.526-0.644mg/l and 0.24 – 1.517 mg/l respectively (**Table. 1**).

**Fig. 9:**
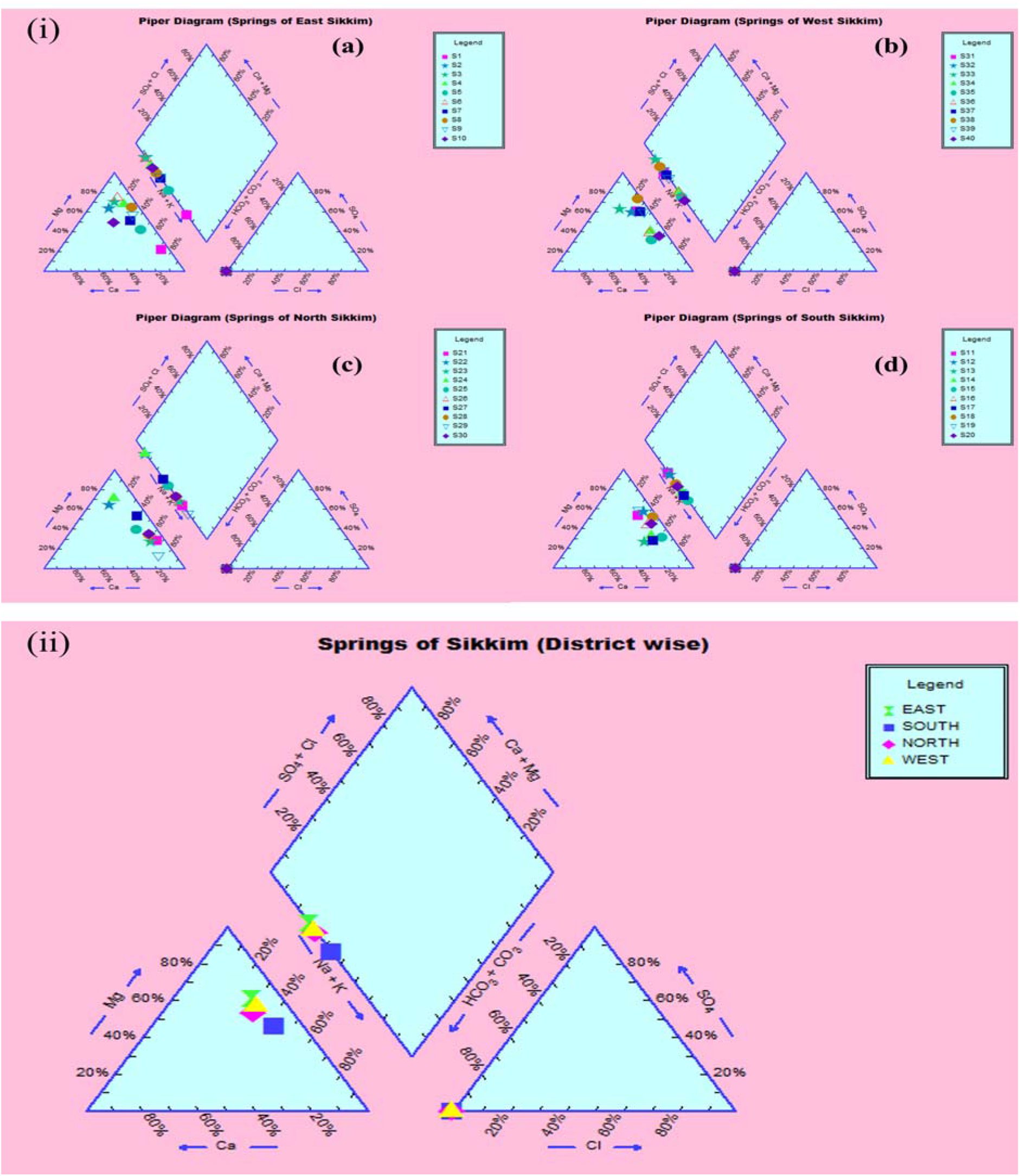
Piper analysis showed waters from all the four districts were Mg-HCO^3-^. (i) District wise classification (a – E, b – W, c- N and d–S) (ii) Collective classification.

Pairwise Pearson correlation analysis between different water quality parameters of the four districts showed interesting correlations. The correlation coefficient (r) determined between parameters Temperature, TDS, Turbidity, EC, TH, Alkalinity, pH are shown in **Fig. 10**. The correlation matrix showed positive correlation between EC and TDS (r = 0.998/1.000, p<0.05) while negative correlation between pH and TDS (r = −0.139/1.000) (**Supplementary File S3 & Fig. 8**). A strong correlation was also observed between Alkalinity and Turbidity (r = 0.993/1.00, p<0.05). While pH, TH and Alkalinity were found to be negatively correlated. The correlation coefficient (r) between alkalinity and pH was r = - 0.108/1.000. Similarly, Total hardness was negatively correlated with alkalinity (r = −0.367/1.000), and pH (r = −0.405/1.000) (**Supplementary File:S3**). Correlation analysis between the chemical parameters showed the concentration of Pb and Al are highly correlated with a correlation coefficient (r) = 0.975 /1.000 (p<0.05). A high positive correlation was also found between concentration of Ag and Al (r = 0.948/1.000), Gallium and Barium (r = 0.988/1.00, p<0.05), Sodium and Calcium (r = 0.980/1.000, p<0.05), Chromium and Barium r (Cr and Ba) = 0.996/1.00 (P<0.05), r (Mg and K) = 0.96/1.000 (p<0.05) respectively. A strong correlation were also observed between Potassium and Calcium (K and Ca) r = 0.95/1.0 (p<0.05), Magnesium and Sodium (Mg and Na), r = 0.99/1, p<0.05). A strong negative correlation was found between Li,U, Xe, Al and Ag [r (Li and Xe) = - 0.99/1.000; (p<0.05), r (Li and Ag) = - 0.769/1.000); r (Li and Al) = - 0.880 /1.000 (p<0.05)], U and Ag [r = −0.97/1, p<0.05)], (**Supplementary File S3 & Fig. 10**). Principal component analysis of the major physicochemical parameter showed all the districts were positively correlated. East districts showed strong correlation with South [r = 0.99/1, p<0.05)] and West [r = 0.98/1, p<0.05)] as well as positive correlation with North [r = 0.94/1, p<0.05)]. Along with the East, South district also showed a strong correlation with West [r = 0.99/1, p<0.05) and North district showed a positive correlation with East as well as south [r = 0.93/1, p<0.05] and least positive correlation with West [r= 0.89/1, p<0.05]. Clustering based on the principal components showed a correlative cluster for E, W and S while N district formed a different cluster defining the difference in physicochemical parameters (**Fig. 11**)

**Fig. 10:**
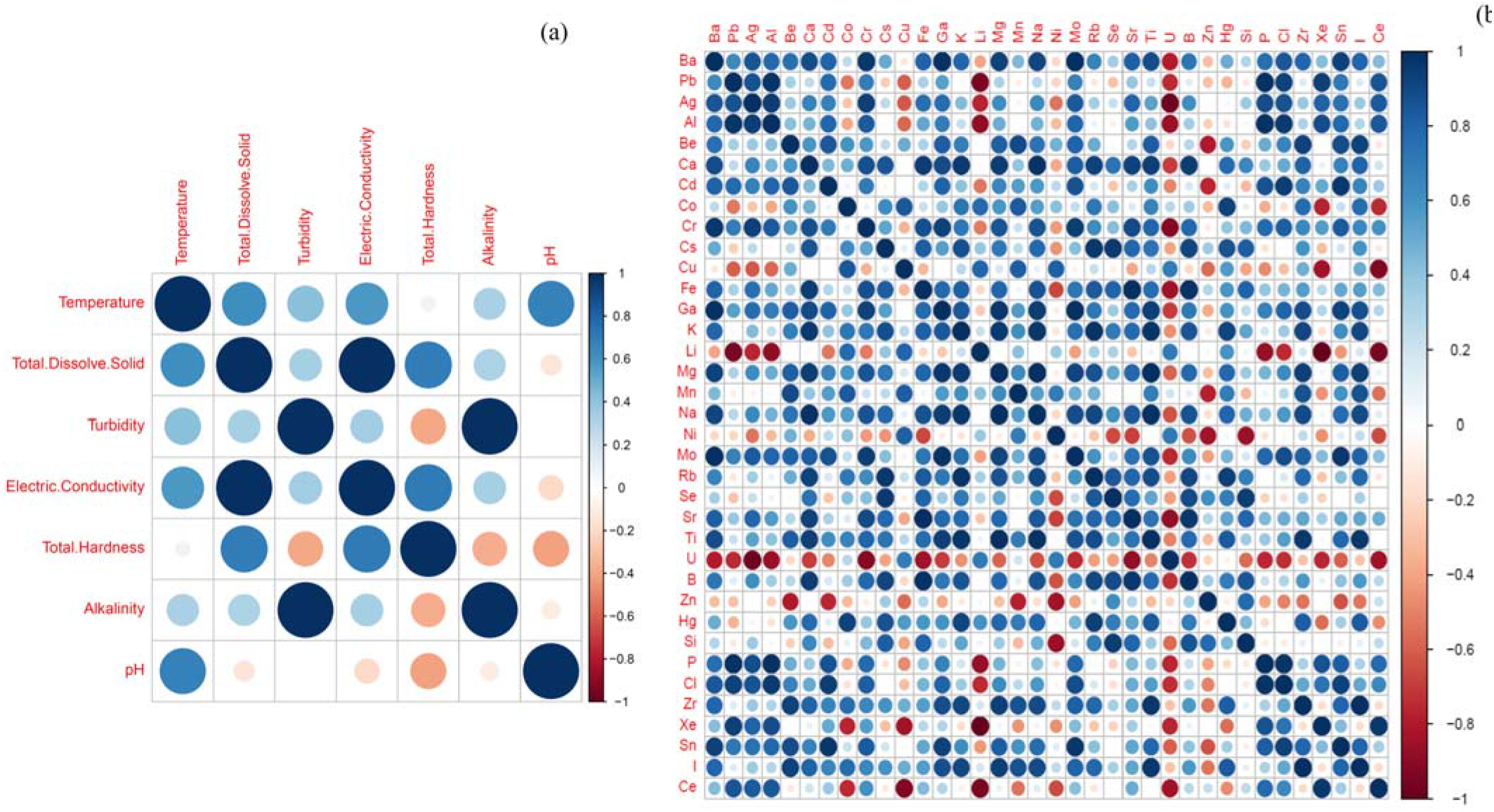
Pearson correlation plot between physicochemical parameters (a) Basic parameters and (b) Chemical elements. The dark blue colors define high positive correlation and dark red color defines high negative correlation.

**Fig. 11:**
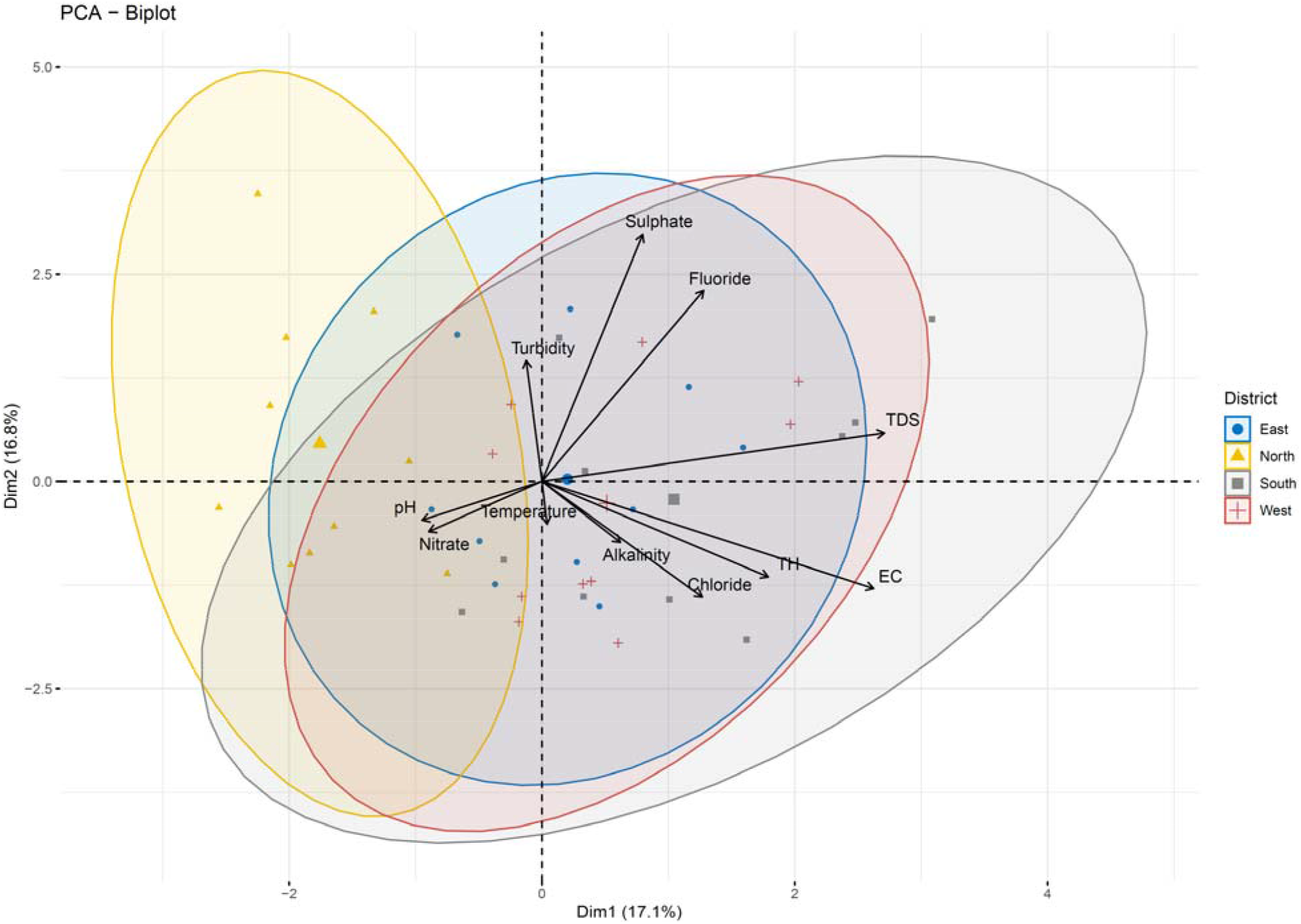
Biplot analysis of the physicochemical parameters across the four districts.

Mean concentration of all the three indicator bacteria in the spring water, community reservoir and household water of four districts were higher than standard WHO guidelines (must not be detectable in per 100ml of water) (**Fig. 12**). The spring water from the East was found to have a total coliform count in the range of 16-44.4 in the three seasons (Rainy, Winter, and Spring) while TC in the rainy season was highest with (64)/ml of the sample. The TC count in the South, North, and West district ranges from 14.9 – 42.8/ml, 11.9 – 31.5/ml and 24.4 – 48.9/ml respectively. Rainy season (July – August) accounted for the highest count in all the districts, East = 44.4, South = 42.8, North = 31.5 and West = 48.9. Community reservoirs were recorded with an astoundingly high number of total coliform bacteria with a highest average mean of 75.6 CFU/ml in the East district in the rainy season followed by South = 71.1/ml, West =67.6 /ml and North = 41.4/ml (Supplementary File: S4). Household water was recorded with the least number of coliform when compared with the spring water and community reservoirs. Household water from the East district had a mean Total coliform concentration of E=13.36, S = 13.52, W= 9.28, and N = 7.2 in the rainy season. Winter season was recorded with the least number of coliforms in all the three sources of water and in all the four districts. The highest coliform count in winter was recorded from community reservoirs of the East district with a mean concentration of 55.2/ml sample. Highest *E. coli* count was recorded from the community reservoirs of the East (13.5 /ml) in the rainy season (July – August) and lowest was recorded from the household water from the North (0.4 /ml) in the winter season. *Enterococcus* was also recorded in high number from community reservoirs in the rainy season. District wise the highest *Enterococcus* was recorded from the East district (5.5 /ml) followed by South: 5.2, West: 2.6 and North:1 respectively (**Fig. 12**). Cluster analysis based on the mean concentration of total coliform, *E. coli* and Enterococcus formed three distinct sub-clusters. South and West district formed one single cluster and which had similarity with East district while North formed the out-group as observed in **Fig. 13**.

**Fig. 12:**
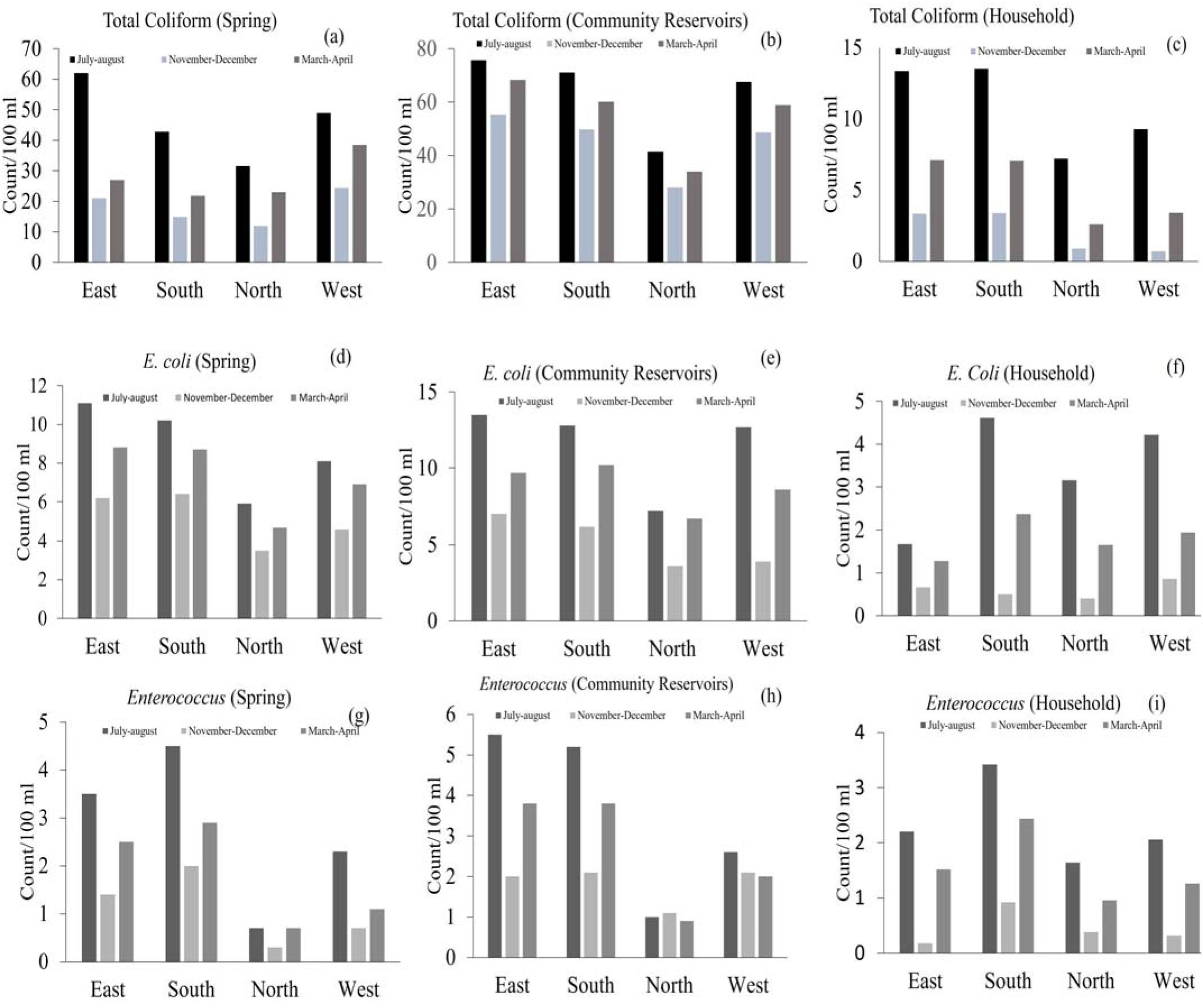
Distribution of Total coliform (TC), *Escherichia coli* (EC) and *Enterococcus* (EN) bacteria in the Springs (SP), Community reservoir tank (CR) and Household water (HH) in the four districts. (a) TC in SP (b) TC in CR, (c) TC in HH, (d) EC in SP, (E) EC in CR, (F) EC in HH, (g) EN in SP, (h) EN in CR and (i) EN in HH.

**Fig. 13:**
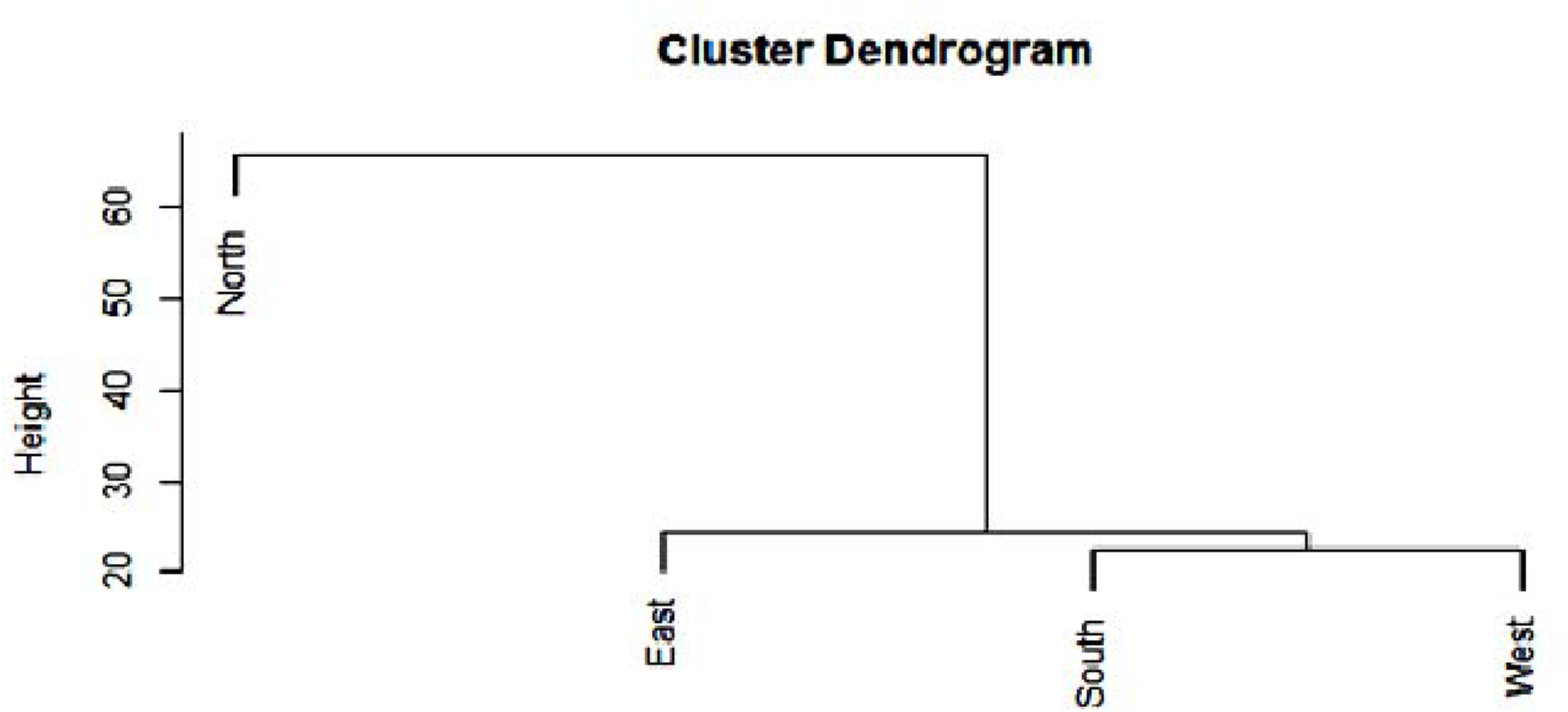
Cluster analysis of the water from E, W, S and N based on the mean concentration of Total coliform, *Escherichia coli* and *Enterococcus*.

## 4. Discussion

Water is essential to life, access to safe and adequate water is important to human health. Providing safe and clean water is a basic norm in developed countries but many people, especially in developing countries, struggle to get access to safe water (Prüss-Ustün et al., 2014) (Alves et al., 2018). Water systems with inadequate quality control and sanitation act as a vehicle for different pathogenic microorganisms which are either originating from feces of human or wildlife. Though huge efforts are put to provide safe drinking water but still, it is estimated that 1.1 billion people are consuming water sources regularly contaminated with fecal microorganisms (Bain et al.,2014; WHO, 2014). Detecting each waterborne pathogen is complex; therefore, a standard reproducible microbial water quality test was developed by the International Organization for Standardization (ISO). In this method, detection of certain groups of bacteria acts as an indicator for the fecal contamination (Saxena et al., 2015). The most widely used indicator groups are *Total coliform, E. coli* and *Enterococci* (Saxena et al., 2015). Though, not only the bacterial pathogens, several physicochemical parameters also determine the water quality of a region. There are standard guidelines by both WHO and BIS for the safe and hygienic water index ((Bureau of Indian Standards, 2012)(World Health Organization, 2017)).

Sikkim, an Eastern Himalayan state with high altitudes as topological features. Majority of the population depends on the naturally occurring spring waters for their day to day use. To maintain the continued supply of spring water, the government of the state had made community reservoirs which are responsible for storing the spring water and supplying to households of an area. But looking to the recent outbreaks of several waterborne diseases around the state has raised a concern regarding the quality of the water consumed by the residents. In this study, we tried to look at the water quality of the four districts of the state with respect to both the physicochemical and microbiological profile. The filtration and purification processes are one of the less common practices among the hill population of the state. Boiling of the water is the one or only common practice used for the sterilization. On average 80% of the population of North and 66% of the East uses boiled water for drinking purpose. Similarly, 56% of South and 60% of West uses the boiled water for drinking purpose. Boiling drinking water gives extensive protection but not completely. There were reports of effective protection against *Vibrio cholera* infections, *Blastocystis*, protozoal infections overall, viral infections overall, and nonspecific diarrheal outcomes by simple boiling method (Cohen and Colford, 2017). The results were correlative with our finding where the household water was recorded with the least number of microbial contaminants where boiling (**Fig. 7**) was applied prior consumption as compared to direct consumption. Only a total of 10% household water was found to be safe to drink, where direct spring water/water supplied from community was used (Coliform count > 1) in comparison to the 33.33% safe water where boiling is done before the drinking and even in rainy season where occurrence of waterborne disease is high (July – August) (**Supplementary File: S4**). The *E. coli* count was also found significantly low in boiling water with 81.81% of household water in East showed *E. coli* count <1/100ml in comparison to 66.66% of direct spring water (Supplementary File: S4). In a research by, Clasen et al., (2008), where he did a 12 weeks of study among 50 households from a rural community in Vietnam, boiling was associated with a 97% reduction in geometric mean of thermotolerant coliforms (TTC) (*p* < 0.001) and despite being high levels of fecal contamination in source water, a 37% reduction in count of total coliform can be observed when compared with the raw water. Even with the benefits of the boiling of water, still, there was the significantly large number of populations of the state who were still using the spring water without any treatments. In East, 20% of the population; West = 10%, S = 30% and North =16% population still uses the spring water directly for drinking. This increases the chances of an immediate health risk as the microbial quality of the water indicates severe fecal contamination.

The community reservoirs which were developed to work as a storage and supply station for the households in village level blocks were the most contaminated ones. The CR from E had the highest TC count in rainy season with a range of 45-93/100ml with a mean of 75.6, which is not acceptable for a drinking water standard. The CR water was also recorded with a high number of *E. coli* and *Enterococcus* too, E. coli (E): 3 −28/100ml (J-A) with a mean of 13.5/100ml and Enterococcus (E):0-13/100ml (J-A) with a mean concentration of 5.5/100ml. This high number of fecal coliforms in water source indicates the severity of the situation and there is a high chance of human or animal fecal contamination somewhere in the source or during transportation of the water from the source to reservoirs. Correlation between the SW, CR and HW showed a significant correlation between the SW and CR (r = 0.90; p< 0.05) indicating the contamination at the spring water source affects the contamination level in community reservoirs. There was also a strong correlation between the CR and HW (r = 0.78) but which was – statistically not significant (p >0.05), thus the contamination at CR may not be the sole reason for the contamination at HW level (**Fig. 14**). The high number of coliform bacteria in the water source is a concern for the state, as increased coliform in the source can become a reason for the proliferation of other opportunistic waterborne pathogens like *Klebsiella, Enterobacter* and *Pseudomonas*. Coliform generally forms biofilm at the site and provides the anchorage and proliferation site for the opportunistic pathogens (Camper et al., 1998; Ridgway et al., 1981; Manz et al., 1993).

**Fig. 14:**
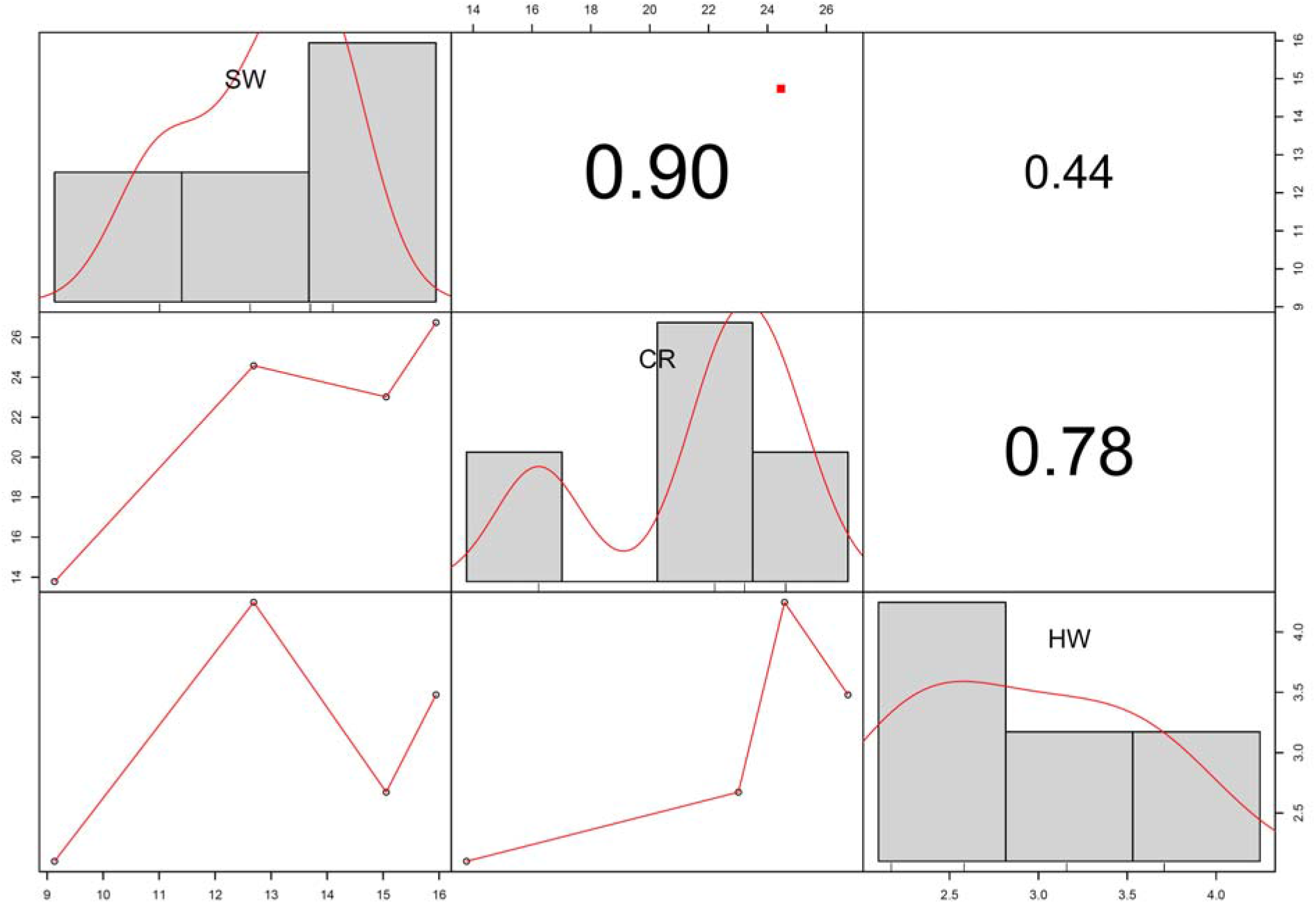
Correlation between the water sources, spring water (SW), community reservoirs (CR) and household water (HW). The red dot above between the SW and CR (r = 0.90) indicate the statistical significance (at p< 0.05).

Water source from the East was highly contaminated with coliform bacteria in comparison to the other state and North is the least contaminated. There was no significant statistical difference among the water sources of four districts (p > 0.05) defining all the four districts were in the alarming zone. The UPGMA cluster analysis divided the water into three subclusters, South and West forming a cluster showing the similar concentration of indicator bacteria while North formed the out grouped with least number of contaminants and East formed one cluster, which is highly contaminated followed by S and W (**Fig. 13**). The severity of water contaminants in the East district is of more concern as it is the main economical or industrial or educational hub for the state with the highest population (45.3%) (Department of Economics, Statistics, 2013). Inversely it also indicates the severe contamination is happening due to the increased animals or human activity which needs to be taken care off sooner than ever. North district has the lowest population in comparison to other districts which may be the reason for the less human contact and less human-made contamination. But this report suggests a serious immediate health risk which could happen until and unless government and public healthcare personal formulate policy structures to protect the important potable water sources of the state from human and animal interventions.

The physicochemical analysis showed most elements were within the standard permissible guidelines, although the presence of few toxic metalloids above the guiding value of WHO, ISI, CPCB and ICMR in the spring water can pose serious health hazards. Elements like Mercury, Cadmium, Selenium, Lead, and Chromium were detected in the spring water of four districts, which are toxic to human (**Table.1**). Mercury was detected in the range of 0.001 – 0.007 mg/l with East district having the highest concentration of 0.007 mg/l. Cadmium, Selenium and chromium were detected in the range of 0.011 – 0.026 mg/l, 0.526 – 0.644 mg/l and 0.249 – 1.517 mg/l. Most of these elements are under the condition of “no relaxation or <0.01-0.05” by Central Pollution Control Board (CPCB) and the World Health Organization (WHO) (Central Pollution Control Board, 2018; Enderlein et al., 1996). Detection of these heavy metals poses a serious concern over the health of the people. International Agency for Research on Cancer (IARC) suggested that inorganic As and Cd are classified as a carcinogen. Cd is related to cancer skin damage, kidney damage, and heart disease; Sb causes high blood cholesterol; Pb is related to anemia; Hg causes kidney and liver damage.; Cu is reported to be associated with gastrointestinal disease (QA and MS, 2016). Physical parameters like pH, Turbidity, TDS, and Alkalinity etc were recorded with the range of standard permissible value as recommended by WHO, CPCB and ICMR. Pearson correlation analysis showed a high positive correlation between the concentration of TDS, and EC (r = 0.998/1.000), Turbidity and Alkalinity (r = 0.993/ 1.000). While TDS, pH, Turbidity, TH and Alkalinity were found negatively correlated with coefficient of correlations as, r (TDS & pH) = - 0.139/1.000; r (Turbidity & TH) = - 0.388/ 1.000; r (TH and pH) = - 0.405/ 1.000; r (TH & Alkalinity) = - 0.367/1.000 and r (Alkalinity & pH) = - 0.108/1.000 respectively (**Supplementary File: S2**). Correlation between inorganic elements showed a strong positive correlation between the concentration of Pb and Al with correlation coefficient of (r) = 0.975 /1.000, Ag and Al with r = 0.948/1.000, r (Na and Ca) = 0.980/1.000, r (Na and Ba) = 0.918/1.00, and r (Mg and Ca) = 0.947/1.000 respectively. A negative correlation was observed between Li, Pb, Al and Ag [r (Li and Pb) = - 0.942/1.000; r (Li and Ag) = - 0.769/1.000); r (Li and Al) = - 0.880 / 1.000 (**Supplementary File: S2 & Fig. 8**). The presence of an excessive amount of physical, chemical and biological parameters in the water sources leads to the effects the human health. As discussed in the result, some of the physical parameters like pH, electric conductivity, turbidity and alkalinity and some chemical parameters like Iron, Magnesium, Copper, Mercury, Cadmium, Selenium and Lead were above the WHO limits. The higher concentration of chemicals and deviation from various physicochemical and biological parameters are a health hazard. Along with the physicochemical parameters, biological parameters like Total coliform, *E. coli* and *Enterococcus* were above the WHO limits (**Fig. 12**) (World Health Organization, 2017). The presence of coliform bacteria in high concentration in the water samples both in spring as well as in community reservoir indicates that water was contaminated with warm-blooded animal’s fecal matter like humans or wild animals and not safe for drinking. This urges the need for proper protection of the major water sources at the supply level or at the source of origination to minimize the human and animal contact. The household water samples were also contaminated with fecal coliform bacteria above the permissible standard which indicates severity and necessity of the awareness about different sterilization and filtration process at the household level. The seasonal study showed that rainy season was a major contributory factor in microbial contamination and which needs much attention. In the rainy season, the water should be properly chlorinated and sterilized at the community level before supplying to the households.

## Conclusion

In Sikkim, as the community considers spring as a holy and safe for drinking and household purpose, but in this study, it was found that most of the spring and community reservoir water comes under the intermediate health risk level category as per WHO recommendations 2018. The physicochemical parameters analysis revealed that almost all parameters were not within the safe range of recent WHO, ISI, ICMR and CPCB guidelines, 2017. Most of the water sources were contaminated with traces of heavy metals which indicates an existing health hazard. The microbial parameters were also higher than WHO standard guidelines in the water sample of springs and community reservoir water samples. The community reservoir water samples were found to be more contaminated than spring water. Interestingly, some of the household water samples were safe as per the WHO guidelines in terms of microbial concentration but still, a majority was contaminated well above the permissible limits. The springs of East and South Sikkim were more contaminated than the other districts and least microbial load were found in the Northern district. This study revealed that the water sources are not safe for drinking and more needs to be done for the primary treatment of Springwater such as adequate chlorination of community reservoir water samples before supplying to households etc. The community needs to be educated about general hygiene, different processes/ methods available for household-level water treatment to make it safer for the consumption and alleviate the risks associated with the use of existing water supply.

## Acknowledgment

The authors wish to thank and extend their sincere gratitude to all the communities of the rural areas of Sikkim, India for their cooperation and active support of the study. The authors also wish to thank the State Institute of Rural Development for their helping hand during sample collection and survey work. The authors are also would like to thanks to Gram Panchayat Unit (GPU) for continuous help and support during survey and sample collection. The authors also like to thank all the faculty members and non-teaching staff of the Department of Microbiology, Sikkim University for their continuous support and help throughout the study.

